# A Tape Spring Steerable Needle Capable of Sharp Turns

**DOI:** 10.1101/2023.05.04.539394

**Authors:** Omar T. Abdoun, Mark Yim

## Abstract

**Objective:** To make steerable needles more effective, researchers have been trying to minimize turning radius, develop mechanics-based models, and simplify control. This paper introduces a novel cable-driven steerable needle that has a 3mm turning radius based on tape spring mechanics, which sets a new minimum turn radius in stiffness-matched tissue models. Methods: We characterize the turn radius and the forces that affect control and performance and create predictive models to estimate required insertion forces and maximum insertion depth. Finally, we demonstrate the performance of a task outside the capabilities of a conventional needle. Results: Minimal force is required to maintain bends, allowing surrounding tissue to fix them in place, and minimal energy is required to propagate bends, allowing the device to navigate easily through various tissue phantoms. The turn radius of the device is independent of surrounding tissue stiffness, making for simple and precise control. We show that all aspects of performance depend on minimizing the tip cutting force. Under ultrasound guidance, we successfully navigate into and then follow a deep blood vessel model at a steep angle of approach. Conclusion: This design allows the system to accurately control the direction of the device while maintaining a smaller turn radius than other steerable needles, providing the potential to broaden access to challenging targets in patients.

## I. INTRODUCTION

Steerable needle technology has the potential to change how biopsies, ablations, and targeted drug deliveries are performed. Physicians commonly use straight needles and probes to navigate to areas in human tissue accessible by straight trajectories. Steerable needles could navigate around sensitive structures or obstacles, or be adjusted in real time during the procedure, increasing the accuracy and precision of treatment location [1]. Steerable needles have been receiving attention from the robotics research community on account of these promises, including several robotic steerable needle0020survey papers [2], [3]. These needles often involve complex non-linear mechanisms and interactions with tissue requiring robotic control and modeling to steer them [4].

Previous approaches to designing steerable needles are numerous. Flexible needles with asymmetric “bevel-tips” [5] are one type. They create curving paths where asymmetric forces at the bevel tip cause the path of the needle to curve as it penetrates the tissue. To move straight they continuously twist while translating forming a helical path. Unfortunately, these paths are limited to large curvatures (typically many centimeters in minimum turning radius) that will vary in shape as the stiffness of the tissue varies. These larger turning radii limit the reachable space. Other approaches include concentric pre-curved flexible tubes, flexible needle with pre-curved stylets, and flexible needles with an articulated tip, among others summarized by da Viega et al. [6]. The concentric tube approach forms complex paths by exploiting the natural curvature of each tube, controlling the interaction of the tubes through rotation and extensions. These devices also have a large turn radius. Steering these devices in tissue is non-intuitive, computationally intensive, and thus difficult without robotic control [7], [8].

The only steerable needle currently commercially available, the Morrison needle, consists of a stainless-steel flexible needle that surrounds a pre-curved multi-core inner stylet. The inner stylet is held in a straight orientation until a control lever is turned, moving one core past the other and allowing the pre-curved portion to assume its curved orientation. The Morrison needle has a minimum turn radius of 80mm with a maximum of 10mm deflection. In the process of curving, it can stress and tear the adjacent tissue along the length of the curving needle which both complicates accuracy due tissue inhomogeneity and damages tissue [9]. A research group that has achieved a 6.9mm turn radius uses a mechanism, creating and following tissue fractures, that has the disadvantage of tearing through surrounding tissue in order to navigate while also being subject to errors from changing tissue mechanical properties [10].

Development of steerable needles requires improvement in four areas to achieve wide application in the real world [19]. They are:

1. mechanics-based modeling,
2. planning in 3D with uncertainty,
3. human in the loop control, and
4. minimizing radius of curvature.

The first three items involve the accuracy of steering the needle and are primarily complicated by the effects of different tissue mechanical properties on the motion of the needle. A needle that imparts minimal stresses on the tissue as it traverses through it will also be less sensitive to tissue stiffness variation. In the ideal case, decoupling of the turn radius of the needle from the tissue properties means 1) a very simple model can be used to predict and control motion, 2) the planning uncertainty is greatly reduced, 3) humans (e.g., surgeons) can control the needle without complex machinery as it is intuitive.

An ideal steerable needle is stiff enough to penetrate the tissue and maintain control of where it goes, yet also flexible when making tight turns. A tape spring is a long, thin, uniformly curved sheet of material that can be reversibly bent without plastically deforming. This tape-spring is made of a stiff but relatively elastic material, such as stainless steel, with a longitudinal curve around the long axis of the strip.

This paper presents design and analysis of tape springs as the shaft of a steerable needle that can create sharp turns that result in reduced stress on surrounding tissue promising the reduction of tissue damage. Lower stress also means more independence of the path geometry from tissue properties thus easing modeling requirements which is an issue for other steerable needles [20].

This paper is organized as follows: Section II details the tape spring used as a steerable needle. Section III describes the manufacturing of the needles as well as experiments that characterize their behavior and performance. Section IV presents the results and discussion of the experiments. Finally, we discuss conclusions and future work.

## II. DEVICE DESIGN AND PRINCIPLE OF OPERATION

Based on the promise of needle access inside the body for a large range of medical uses, our steerable needle has four performative goals:

1. Minimize turning radius.
2. Minimize tissue damage.
3. Penetrate to depths useful for treatment.
4. Minimize cost and complexity while maintaining ease of use by physicians.

A tape spring is a long, thin, uniformly curved sheet of material (typically spring steel or a composite) that can be reversibly bent without plastically deforming. This tape-spring is made of a stiff but relatively elastic material, such as stainless steel, with a longitudinal curve around the long axis of the strip.

The most common applications for tape springs, which leverage the ability of such a structure to reversibly bend, are in measuring tapes, solid state hinges, and deployable space structures [21], [22]. Critically, tape springs are stiff when straight, as the transverse curve greatly increases the second moment of inertia, but it can be buckled when a bending moment is applied. Once buckled, the transverse curvature of beam is flattened at the bend, having a much smaller second moment of inertia, which leads to bends maintained with minimal force and small turning radius.

We leverage three properties of tape springs in the design of a steerable needle: 1) reversible elastic buckling, 2) axial stiffness higher than bending stiffness, 3) minimal energy required to propagate a bend from its initial location to an adjacent one. This allows the tip of the device to be actuated in a desired direction and for the shaft of the device to follow the path of the needle tip as it is pushed further into the tissue. The tip can be actuated to various degrees, either right or left, and a plane of motion can be initially selected by rotating the needle. Figure 1 shows the general design of the tape spring steerable needle in both a bent and unbent state. As in the Morrison needle, if an out-of-plane motion is required, the needle must be retracted and reinserted without curvature.

**Fig. 1:**
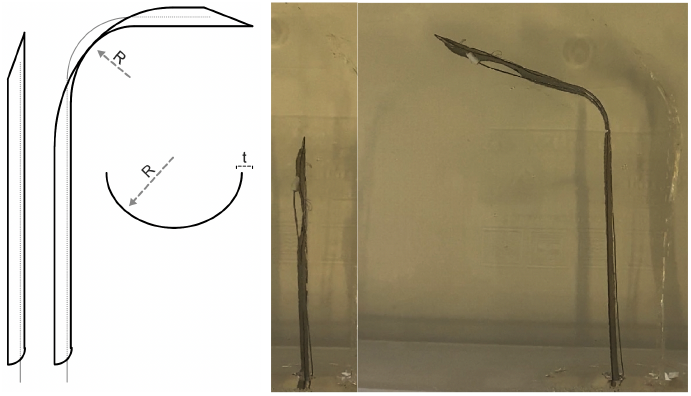
From left to right: a) A schematic of a straight, unbent tape spring needle. b) A schematic of a tape spring needle with an induced bend and *R* as the turn radius. c) A schematic of a head-on view of an unbent needle showing *t*, the thickness of the sheet, and *R*, the transverse radius of curvature of the tape spring. d) A straight, unbent tape spring needle. e) A tape spring needle with an induced bend.

### A. Reversible Elastic Buckling and The Relationship to Axial Stiffness and Bending Moment

The main characteristic of the tape spring as an underlying mechanism for a steerable device is on-demand reversible buckling. Pellegrino et al. [21], [22] described these mechanics. During the process of bending a tape spring, the maximum stress is found at the midpoint of the arc formed by the transverse cross-section of the segment bent, and the longitudinal radius of curvature of the bend is approximately equal to the transverse radius of curvature of the unbent tape spring. Therefore, the minimum turning radius for a tape spring-based steerable needle is approximately equal to its starting transverse radius of curvature.

In designing a tape spring device, material and geometric considerations must be made in order to ensure that the point of maximal stress does not experience stresses that exceed the yield stress of the beam and are described by Equation 1:

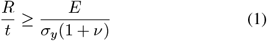

where *R* is the transverse radius of curvature of the unbent tape spring, *t* is the thickness of the material, and *E, σ*_*y*_, and *ν* are the Young’s modulus, yield stress, and Poisson’s ratio of the material, respectively. This places a lower limit on the curvature, and thus, a lower limit on the radius of curvature for a bend made by that tape spring for a given *R/t*.

Our design allows the user to precisely control the direction and angle of bending of the tip, which then controls the path of the needle through tissue. Cables that run along the shaft of the device, from the tip of the needle to the user, are used to bend the device in the desired direction, in a manner similar to how some catheters are steered. Unlike catheters which require an existing channel such as a vein, our device cuts tissue to make its own channel.

When enough tension is applied to one of the cables, the restoring forces from the stiffness of the structure are overcome and the needle tip locally bends towards the side that the cable is attached to at the tip. The minimum moment required for this behavior depends on the geometric parameters of the tape spring, the Young’s modulus, as well as the direction of bending. Equal-sense bending is defined as bending in which the initial transverse center of curvature is on the same side of the tape spring curve, whereas opposite-sense bending occurs when the newly formed longitudinal center of curvature is on the opposite side [21], [22]. This is shown in Equation 2 for equal-sense and opposite sense bending, respectively:

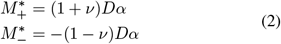

where *D* is defined as: *D* = (*Et*^3^)*/*12(1 − *ν*^2^). From this, we can see the critical buckling moment is asymmetric. Notably, for a *ν* of 0.3, which is the approximate *ν* for stainless steels, 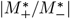 is 1.9. This means that a stainless steel tape spring requires nearly double the force to undergo equal-sense bending than opposite-sense bending. If necessary, this can be mitigated by incorporating an asymmetric transmission, giving a larger lever arm to the side requiring the larger moment.

### B. Bend Stability and Bend Propagation

As a tape spring moves forward into the tissue, a bend in the path propagates down the needle such that the body of the needle follows the path set by the tip. While the moment required to initiate the bend may be large, the moment required to sustain and propagate a bend can be negligible with the right materials [21]. The strain energy required to bend the straight part into the beginning of the arc is recovered by the unbending at the end of the arc. This bend propagation is pictured in Figure 2. Internal energy losses from a turn come from the damping factor of the tape spring’s comprising material(s). This factor describes the energy lost in the stress-strain hysteresis curve of a material as it bends and unbends. While some materials, such most plastics and nickel–titanium alloy, are highly elastic and often used in medical devices, the high damping factors lead to large bend stresses during bend propagation. Metals such as stainless steel, can have low damping factors.

**Fig. 2:**
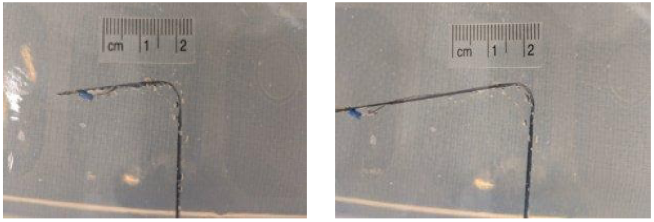
Turns performed by the steerable needle in 7kPa gel. The steerable needle is advanced into the gel with a *∼*3mm radius bend and the bend propagates down the needle. Note the arc length remains the same.

It is important to minimize these stresses as the forces that maintain this bend come from stresses on the local tissue. So, these smaller moments result in smaller stresses on the tissue which reduces tears on increased damage.

### C. Forces Acting on Needle in Motion

Using the terminology similar to previous needle insertion studies [23], the needle-tissue interaction forces can be decomposed into four parts: *F*_insertion_, *F*_cutting_, *F*_friction_ and *F*_stiffness_, respectively. The force outside the tissue pushing the needle into the tissue is the insertion force, *F*_insertion_. The cutting force, *F*_cutting_ is the force encountered at the tip of the needle that is required to cut through the oncoming tissue/gel, and depends on tip geometry, tissue/gel material properties, and needle velocity. The friction force *F*_friction_ is the shear force between the surrounding gel/tissue and the body of the needle and acting along the length of the needle.

Okamura [23] present this equation relating the four terms.

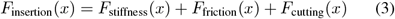

Here the friction is the Karnopp model, dominated by viscous friction which is proportional to the surface area parallel to motion. In addition, the cutting force is considered to be linear with velocity [24] [25]. The reader is referred to Okamura [23] for details.

In [23], the stiffness term primarily relates to the deflection of a tissue as a needle starts to puncture then transitions to a steady state as the needle enters at a constant velocity. In our case, where we are primarily interested in the effects of bending, we will examine just the steady state case after the needle has punctured tissue, which is the condition of interest where turns may occur.

So, a needle entering without turns in the steady state will not have a stiffness component. When modeling just this portion, we propose that a simplified model has both friction and cutting force proportional to velocity.

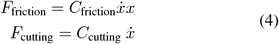

yielding an equation that is more easily experimentally confirmed:

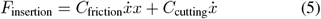

Note that both terms on the right side are proportional to velocity 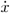. It will be more convenient to refer to a *normalized* force, 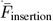 which is 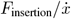.

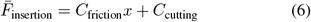

One may question whether a bend in the path results in different equations. Certainly the force balance may be different as the cutting force is no longer aligned with the insertion force. To answer this, we can examine the forces on a needle with a large bend as in Figure 3.

**Fig. 3:**
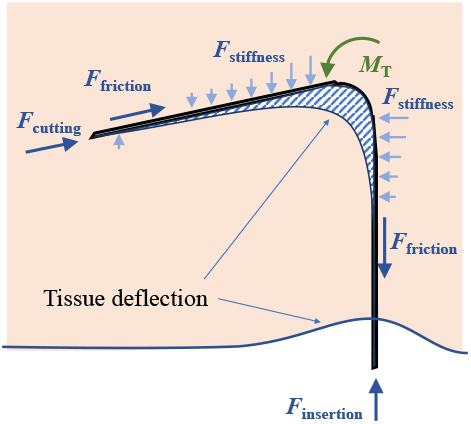
Diagram of external forces acting on a tape spring based steerable needle. Note that tissue deflection is exaggerated for visualization purposes.

Here, the forces that result from the deflection of the tissue, the stiffness force, *F*_stiffness_, is responsible both for the deflection at the entry to the tissue, as well as the reaction forces that hold a bend in the tissue. In practice, what we have observed is that as a bend is established, initially, the amount of tissue that deflects (as shown in the hatched region of the figure) grows, reaching some steady state where deflection stops even as the needle continues at constant speed which is consistent with Okamura’s model [23].

When in this steady state, we can use the principle of virtual work to analyze forces. Here the portions where *F*_stiffness_ applies have only motions perpendicular to the force (the needle slides without any increase in deformation). So, using the principle of virtual work, only the friction, insertion, and cutting forces have non-zero velocities, and thus Equations 5 and 6 hold for bending needles as well.

For analyzing the effect of the sharp bends capable by our needle when not in steady state, it is convenient to consider the needle with a single bend as having two rigid bodies connected by a hinge where the bend would be. There is a proximal component and a distal component, as shown in Figure 4, where forces are labeled with “*D*” for distal and “*P* “ for proximal. With this simplification, we can sum the forces and moments on each rigid body. Looking at the distal body, we note that the forces can be decomposed into two directions; those inline with the long axis (*F*_cutting_, *f*_friction-D_, *f*_Da_) and those perpendicular (*f*_stiffness-D_, *F*_tip_, *f*_Dn_), where labeling with *a* refers to axial and *n* to non-axial.

**Fig. 4:**
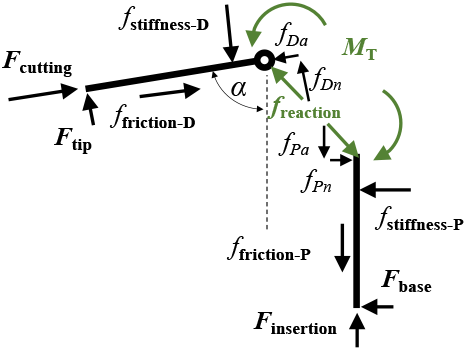
A simplified model where the bent portion is replaced by a hinge and the other beams are rigid bodies.

Summing the forces in axial and non-axial directions on each rigid body yields:

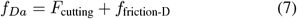

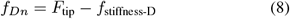

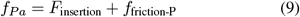

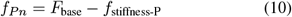

The reaction forces on each component must sum to zero:

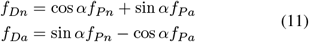

If we also assume negligible internal moment forces at the hinge (as discussed in Section II-B), we can ignore the moment balance and all of the non-axial forces must be zero *F*_base_ = *F*_tip_ = 0 which is reasonable from observed tests.

To interpret these equations it may be most intuitive to examine two extremes, when *α* = 0 and when *α* = *π/*2. In the first case, the needle is straight, so we end up with *F*_cutting_ + *f*_friction-D_ = *f*_*Da*_ = *f*_*Pa*_ = *F*_insertion_ −*f*_friction-P_ and all other forces equal to 0. In the right angle bend *α* = *π/*2 case, combining Equations 7 and 11 we have:

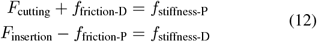

but from Equations 8, 10 and *α* = *π/*2, *F*_tip_ = *F*_base_ = 0 we have again:

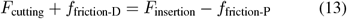

If friction is small, the insertion force becomes equal to the cutting force, which is also equal to each of the stiffness force components. This is important in that the tissue damage from a bend is likely to be a function of the stiffness force. Thus, to minimize damage in this low friction case, we should minimize cutting force.

Even if the friction is non-negligible, it is likely to still be a monotonic function of the insertion force. For example, if we use a model where the friction is linear with a normal force, the normal force for *F*_friction-D_ comes from *F*_stiffness-D_. However, this force is a function of *F*_insertion_ −*F*_friction-P_, and *F*_friction-P_ is linear with its normal force, which scales with *F*_cutting_ and is thus linear with *F*_insertion_, as well.

So, even where friction is significant, minimizing the cutting force will minimize *F*_stiffness_, thus minimizing tissue damage. There are several factors that affect *F*_cutting_ including the sharpness of the needle tip, the bevel angle, and potentially the needle diameter [20].

### D. Tape Spring Buckling

One factor that affects the accessibility to different areas in the body is the maximum insertion length of a needle. When the needles are thin enough to minimize damage and allow for turning to occur, we observed that the deeper the needle goes, larger forces on the needle are required to push the needle, (*F*_insertion_) that ultimately caused the proximal straight portions of the needle to buckle instead of progressing further at the tip. This depth for slender beams under axial compressive loads are normally limited by Euler column buckling.

While Euler buckling and the critical buckling load of a column in space is well understood, the effects of having a compliant material, like surrounding tissue, is less so. Our situation is more similar to piles confined by soils which is a topic that has been studied in civil engineering. Here we adapt the concepts of “Winkler Springs” [26] on confinement and how skin friction is used in pile modeling to build a model of tape spring steerable needle buckling in tissue. A simple factor *S* applies to a scaled Euler buckling load equation:

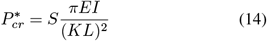

where *S* is the confinement multiplier, *E* is the material Young’s modulus, *I* is the area moment of inertia, *L* is the length of the needle, and *K* is the effective length factor, which is 2 in this case for the fixed-free end-condition.

## III. EXPERIMENTS

The following experiments evaluate the tape spring approach and validate the model presented. In addition, we describe a realistic application (intravenous needle placement) to demonstrate a potential case.

### A. Needle Manufacturing

Strips of steel were cut by hand and by water jet from sheets of 50um thick 301 stainless steel for the 8mm needles and 40um thick sheets of 1095 steel for the 3mm needles. The patterns cut into the strip include features for bend localization, a point at the tip of the needle, and holes along the shaft of the needle for cable routing. Pressing the cut strips into an aluminum mold with a cylindrical cutout imparts the required transverse curvature. Heat treating the strips reduces spring-back, so the curvature remains in the strip after removal from the mold. The tape springs that were tested fulfilled the criterion of the value of *R/t* being greater than that of *E/*(*σ*_*y*_(1 + *ν*)) (Table II) such that reversible elastic bends could be formed, as shown in Figure 1. Finally, cables are routed through the holes along the shaft and attached at the tip using cyanoacrylate adhesive. Important properties for the tape spring device are shown in Table II taken from [27].

**TABLE I:** Comparison of Various Steerable Needle approaches summarized from da Veiga et al. [6]

| Reference | Design | Diameter(mm) | Turn Radius |
| --- | --- | --- | --- |
| [11], [12] | Notched steerable needle | 3.2 | 171 |
| [13], [14] | Concentric tube | - | 72 |
| [15] | Backbone tendon driven | 26 | 76 |
| [16], [17] | Backbone, tendon driven | 10 | 35 |
| [18] | multi-backbone segments | 4.2 | 8.13 |
| This work | Tape Spring | 3 | 3 |

**TABLE II:** Tape Spring Steerable Needle Critical Parameters

| $R$ [mm] | Material | $t$[m] | $E$[GPa] | $\sigma_y$ [MPa] | $\nu$ |
| --- | --- | --- | --- | --- | --- |
| 8.4 | 301 stainless | 5.08E-5 | 193 | 931 | 0.3 |
| 3.0 | 1095 steel | 3.81E-5 | 205 | 800 | 0.3 |

### B. Needle Turn Radius Characterization

While tape spring bends have been characterized for open air or space applications [21], the effects of surrounding material on tape spring bends have not. We present a series of benchtop tests to quantify the turn radius of tape spring-based steerable needles in gels of varying stiffnesses using Knox™Porcine gelatin for gel phantom models at different concentrations. The approximate values of *E* for the tested gels are 7kPa, 13kPa, 36kPa, and 76kPa, obtained by using the method in Henderson et al. [28]. All gels were used immediately after 24 hours of refrigeration at 4°C.

Each trial was performed by inserting our needle prototypes into each tissue phantom and introducing a single 90° bend by applying tension to the steering cable to create a bend at the tip of the needle. Fiji-ImageJ computer vision software measured this turn radius from photographs of each trial.

### C. Steerable Needle Force Characterization

Six sets of experiments measure *F*_insertion_ directly under different conditions so we can derive *F*_friction_, *F*_cutting_, and *F*_stiffness_ components. In all three sets, a steerable needle mounted with its tip facing upwards sits on a digital scale such that any downward force acting on the needle is measured. This scale measures the force as gel constructs of both 7kPA and 13kPa lower onto the needle. A reference tape measure next to the steerable needle at the same distance away from the recording device captures and correlates displacement with the measured forces. Visual markers drawn onto the gel measure insertion length and insertion velocity. Table III summarizes the six sets of trials performed where we varied gel stiffness, insertion length, and needle trajectory.

**TABLE III:** Trial Design

| Trial Set | Gel Stiffness | Insertion Length | Straight vs 90° Turn |
| --- | --- | --- | --- |
| 1 | 7kPa | Constant | Straight |
| 2 | 7kPa | Variable | Straight |
| 3 | 7kPa | Variable | Turn |
| 4 | 13kPa | Constant | Straight |
| 5 | 13kPa | Variable | Straight |
| 6 | 13kPa | Variable | Turn |

In the **constant** insertion length trial (sets 1 and 4), the steerable needle passes straight through a gel construct such that the needle tip extends outside the gel at all times (*F*_cutting_ is 0 and the insertion length is constant). In the **variable** length trial (sets 2, 3, 5 and 6) the needle starts outside the gel and is inserted progressively into the gel. The **straight** trial (sets 1, 2, 4 and 5) proceed without bends, whereas **turn** sets include an approximately 90° bend in the middle of the gel.

We use methods for force analysis similar to Okamura [23] except simplified so we can analyze forces from turns. Since it is more convenient to use velocity normalized force 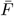 we divide each measured force by the instantaneous velocity determined by dividing the change in position between sequential video frames by the time elapsed.

#### 1) Cutting force

The initial 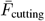 is derived from 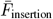 during **variable** trials (sets 2 and 5) at the point where the needle first inserts into the gels. Since the needle is not yet in the tissue, 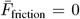, so 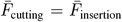 (from Equation 6) once in steady state.

#### 2) Friction Force

The friction model accuracy can be measured in two steps from Equation 5. The first step uses the insertion force from the constant length sets. The measured 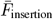 represents the 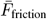 of a fixed amount of gel (fixed *x*) that the needle passes through. Since the needle is straight and the tip protrudes past the gel, the only remaining force must be from friction. With constant *x*, the linear relationship with velocity is apparent and we find *C*_friction_. The second step verifies the Equation 6 is linear with *x* beyond the one measured point in the previous step.

#### 3) Turning Reaction Forces

The 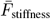 term can be de-rived from Equation 12 where *α* = *π/*2. Since we can determine 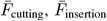, and the friction terms, we are left with the stiffness terms. The moment produced at the bend, *M*_*T*_, is the moment required to sustain a bend that has already been made. That is the residual moment in the buckled beam. In addition, this includes the moment required to propagate a bend through the needle. Note that once beams have buckled they have small *M*_*T*_ and by choosing the right materials, the propagation moments are also very small.

The moment produced at the bend, *M*_*T*_, in practice, acts as a distributed force along the outside of the needle applied to the surrounding tissue and is easily sustained by gel or tissue. *M*_*T*_ was indirectly measured by measuring the force required to sustain a bend that has already been made in the tape spring needle in air where no cutting force or friction force is present. The output force was measured using a digital scale that the tip of the steerable needle was attached to after the needle was bent. The product of distance from the bend to the needle tip and the output force was used to calculate an *M*_*T*_ value. Various distances were used to calculate a mean *M*_*T*_.

### D. Tape Spring Buckling with Confinement from Surrounding Gel

To better understand the limitations of buckling and whether the civil engineering Winkler Springs model applies, a test was designed to measure critical buckling load both in air and surrounded by a gel of known stiffness.

The experiment uses a 102mm diameter pipe that encloses 100mm sections of tape spring used in tape measures. Load cells collect force data. An external frame with a vertical degree of freedom applies an axial load to tape spring segments until they buckle.

### E. Ultrasound Guided IV as a Steerable Needle Application

A realistic application of a steerable needle is intravenous (IV) needle placement. Ultrasound guided IV placement is a common procedure performed in order to place catheters in difficult to reach veins. Even with ultrasound guidance, the depth of a vein limits the angle of approach possible with a conventional needle and increases the risk of “through and through” puncture [29].

The ultrasound guided IV placement trials use Knox™ porcine gelatin at 13 kPa with a target circular channel 10mm in diameter at a depth of 45mm. Manual attempts to navigate into the channel using ultrasound image guidance with both a straight needle and a steerable needle at an approximate angle of approach of 90°. This very steep angle of approach was chosen to demonstrate the maneuverability of the tape spring needle relative to a conventional one. In five trials, the needles were manually advanced until the tip was at the superior aspect of the channel, at which point we attempted to maneuver the needle into the vessel phantom. A successful trial was recorded if the needle could be navigated into the channel without puncturing the bottom of the channel or ripping through the surrounding gel. A medical student trained in ultrasound guided IV placement performed the trials.

## IV. RESULTS AND DISCUSSION

Here we examine two unique aspects of the tape spring needle, the tight turning radius and the independence of steering from tissue properties. We also discuss its limitations.

### A. Steerable Needle Turn Radius Characterization

A survey of interventional radiologists found that the minimal allowable placement error for a device was 2.7mm and that the mean maximal encountered error was 5.3mm. This survey also found that 95% of respondents wanted increased manipulability of their steerable needles and that 93% viewed steerable needles as a potential value-add in their practices [30]. The turning radius enhances the ability to accurately steer a needle as a smaller turning radius is required to adjust offset errors especially as the needle tip approaches the target.

The minimum turn radius of a commercially available steerable needle, is 80mm [9]. The smallest steerable needle turn radius of a steerable needle that we could find by another research group was 6.9mm using a method that requires a flexible needle to follow fractures in the surrounding tissue [10]. This method may not be feasible in real tissues for three reasons: 1) While a goal of any medical intervention is to minimize collateral damage to surrounding tissues, their design requires significant damage to surrounding tissue. 2) While fracture formation can be leveraged in gels due to the tendency of gels to fail because of tensile crack propagation from small defects, it may not be feasible in real tissues due to the higher fibrosity and tensile strength of real tissue as compared to gel models [31]. 3) The design was tested in gel models that are much stiffer than all tissues except bone, tendon, and ligaments, which are not the target tissues for steerable needles [10] [32].

The mean turn radius and standard deviations in mm for all trials are presented in Table IV. Trials included two different transverse radius of curvatures for the needles and various tissue stiffnesses.

**TABLE IV:**
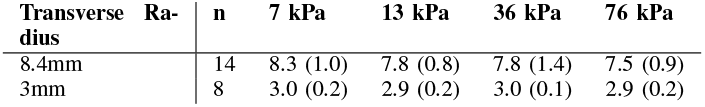
Mean Turn Radius (*σ*) [mm] in different gels

| Transverse Radius | n | 7 kPa | 13 kPa | 36 kPa | 76 kPa |
| --- | --- | --- | --- | --- | --- |
| 8.4mm | 14 | 8.3 (1.0) | 7.8 (0.8) | 7.8 (1.4) | 7.5 (0.9) |
| 3mm | 8 | 3.0 (0.2) | 2.9 (0.2) | 3.0 (0.1) | 2.9 (0.2) |

The overall average turn radii were 7.9mm and 3.0mm for the 8.4mm needles 3mm needles, respectively. The mean turn radius for the 3mm needles was smaller than that of any other steerable needle in current literature or commercially available. In addition, this result agrees with tape spring theory in air where the larger needles 7.9mm is within 10% of the theoretical turn radius of 8.4mm, and in the smaller needles was on average exactly as expected.

Inhomogeneity of tissue is a known factor that makes robotic path planning for steerable needles more difficult and can reduce the accuracy of paths [33]. Table IV also shows that the average for each set of trials falls within one standard deviation of all of the other trials. This implies the turn radius is independent of tissue stiffness for the range 8kpa76kpa. This range spans the range of stiffness’ of organ tissues including, kidney, lens, iris, lung, spleen, thyroid, muscle, and healthy liver to diseased liver with stage IV Fibrosis [32], [34]. The turn radius is independent of tissue stiffness, which suggests the motion of the needle is more predictable than many of the other steerable methods in heterogeneous tissues. In fact, this predictability means that robotic computer-controlled steering is not required and clinicians can use their intuition to guide the needle.

### B. Steerable Needle Force Characterization

Here we will show that the turns in the needle path have surprisingly little effect on the forces. We start with the the linear relationship with velocity (Equations 5) (Figure 5) and the linear relationship with insertion depth. Figure 6 shows the relationship between normalized insertion force and insertion length for all trials. Next, from *f*_cutting_ which should be independent of turns we can derive *f*_friction_ which we can then analyze under straight and turned conditions and infer *f*_stiffness_.

**Fig. 5:**
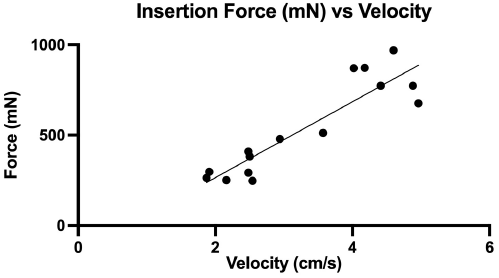
Results of constant length needle insertion force test vs velocity in 13 kPa gel

**Fig. 6:**
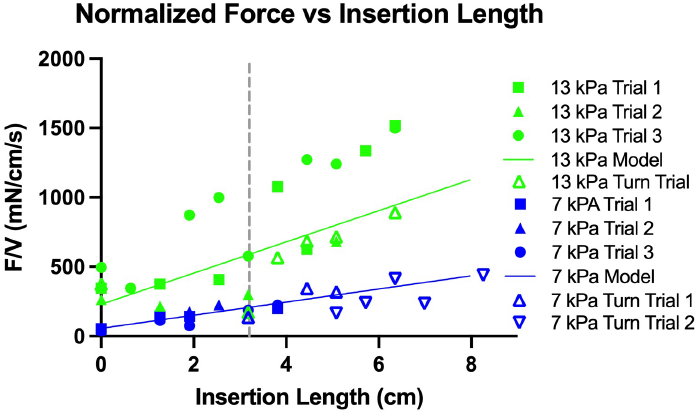
Variable length straight insertion trials show a linear relationship between insertion length and 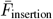.

#### 1) Cutting Force

Linear regression confirms the linearity relationship with with the 7kPa gel having mean *C*_cutting_ = 56.8*mN/cm/s* and 13kPa having mean *C*_cutting_ = 230*mN/cm/s*.

#### 2) Friction Force

While Okamura et al., uses a Karnopp model for friction that is dependent on velocity [23], Fukushima et al., has a model that is not [24], though Fukushima lumps all velocity dependent forces with cutting forces so they may not be contradictory, except in labeling the force components. In our case, during a bend, the friction and cutting forces are not parallel as the proximal component, *f*_friction-P_ is not colinear due to the bend, so lumping forces together does not make sense.

Table V shows the mean (and standard deviation) *C*_Friction_ estimates using both variable length and constant length methods as well as turn tests.

**TABLE V:** Mean *C* _friction_ (Standard Deviation) 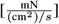

| Gel Stiffness | Constant | Variable | Turn |
| --- | --- | --- | --- |
| 7 kPa | 34.8 (10.1) | 47.3 (7.85) | 42.8 (16.3) |
| 13 kPa | 136.3 (11.3) | 112.5 (44.4) | 122.0 (12.7) |

Friction force was larger in the higher stiffness gel than in the lower stiffness gels. The difference between the *C*_friction_, estimate determined using constant vs variable length was*∼* 26% for the 7kPa trials and *∼*21% for the 13kPa trials.

#### 3) Turning Reaction Forces

For needles with bends in them it is not clear that Equation 6 still holds. From the turned tests, (sets 3 and 6) we can estimate *C*_friction_ assuming the normal forces can be extracted from the simple model from Figure 4 if we set *α* = *π/*2 for a 90° turn. Here the normal force for *f*_friction-P_ would be *F*_cutting_ + *f*_friction-D_ and the normal force for *f*_friction-D_ is *F*_insertion_ −*f*_friction-P_. If the coefficient derived from the turned tests is similar to the one derived from the straight tests, the same friction model could be applied in the case of a turned tape spring due to the relatively small forces required to translate a bend down the body of the tape spring needle.

One might expect a turn in the needle trajectory to affect *F*_Friction_ as new forces normal to the needle surface are introduced. An increase would lead to larger *f*_stiffness-P_ resulting in increased damage as can be inferred from Equation 12. Comparing variable insertion length tests using straight trajectories to trajectories that included a single turn can characterize this potential effect. Surprisingly, the results in Table V show that the measured forces for trajectories with and without turns were very similar. This is also seen in Figure 6 the dotted vertical line indicates the insertion length in the turned tests where the bend occurs. The points for the turned tests stay mostly linear before and after the turn.

The *M*_*T*_ characterization found that the moment required to sustain a bend was less than 0.0015 Nm in all 10 trials. At a distal segment length of 20mm, this translates to 7.5Pa reaction stress of the surrounding gel. Practically, this is a small enough moment that all tissue phantoms used were able to sustain the bend. This distal length was selected because it is approximately equal to the sum of the needle turn radius plus the tip length. As the distal segment length increases when the needle progresses into the tissue, the resultant force is distributed over a larger area.

These results imply the stiffness forces are small compared to the friction and cutting forces, as the addition of a turn did not affect the overall insertion forces significantly.

### C. Insertion length limits due to column buckling

In our buckling experiments, we found that confinement of a tape spring by a surrounding gel increased the critical buckling load allowing deeper penetration than would happen without confinement as expected. Peak buckling force was measured for three different tape springs across different surrounding media to characterize the effect of confinement on peak buckling force. The mean peak buckling force for each specimen in air, 7 kPa gel, and 36 kPa gel can be seen in Figure 7. Each point in Figure 7 represents the average peak buckling force between 15 and 20 trials of each combination of a specimen and surrounding material. Ratios of average peak buckling force in the air to average peak buckling force were used to find S factors 1.36 for 7kPa and 1.89 for 13kPa using linear interpolation.

**Fig. 7:**
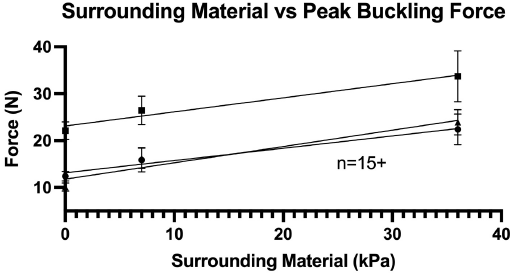
Critical buckling force data across different surrounding media with linear regression.

Measured buckling force at different insertion velocities is compared to two different buckling models in Figure 8. The first, which assumes the confined model of Euler buckling, calculates maximum buckling length from Equations 14 and 5, substituting *F*_insertion_ in for 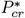 is:

**Fig. 8:**
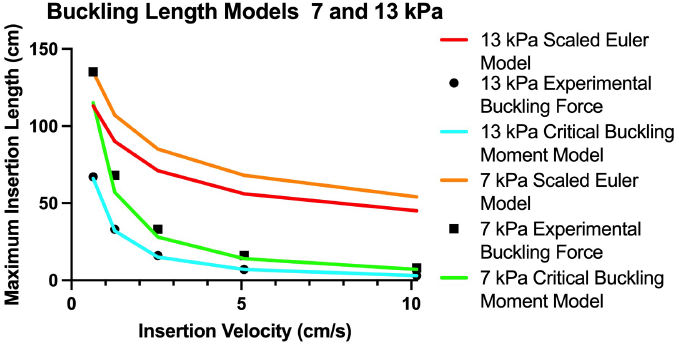
Scaled Euler buckling and critical buckling moment maximum insertion length models.

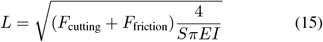

The second model uses the moment buckling equation that describes equal sense bending in tape springs, Equation 2, multiplied by the experimentally derived S factor. 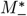 can be deconstructed into a force, *F*_*M*_ and a perpendicular distance, *d*_offset_ as 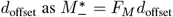. Substituting *F*_insertion_ from Equation 5 for *F*_*M*_ and replacing *x* with *L* in Equation 5, the maximum insertion length, yields:

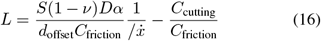

where *d*_offset_ is selected as 1mm to represent a small translation due to the flexible nature of the tape spring steerable needle.

As can be seen in Figure 8, the same sense buckling moment model predicts the buckling length surprisingly well. Since the buckling force predicted by the moment model is smaller for a given velocity and insertion length, even while the Euler-based buckling model may be valid, the beam buckles before the insertion force reaches that level.

It is also clear from the model that velocity is an important factor that affects the maximum insertion length of the tape spring needle. In practical terms, a surgeon using this device could theoretically minimize the chance of placement errors due to buckling by decreasing the insertion velocity. This aspect of the needle design would need to be balanced with the preferences of a surgeon, surrounding tissue mechanical properties, and the time constraints of a given procedure. An optimal set of insertion velocities in a given organ could be a subject of future study. In the case of healthy liver with an estimated tissue stiffness of 7 kPa, the estimated maximum insertion length would be about 30cm at 3 cm/s, as can be seen in Figure 8. If the liver is more fibrosed and has a tissue stiffness of 13kPa, with an insertion velocity of 3 cm/s, the maximum estimated insertion length would drop to about 20cm. This is practically the equivalent effect of almost doubling the insertion velocity in healthy tissue.

### D. Design Considerations

There are a variety of design issues in the current prototype we hope to address in the future.

**The needle size is too large** for most practical medical uses. Our current smallest needle prototypes have a *∼*3mm diameter, which is similar to other steerable needles [6]. But, the target width is 2.67mm or less based on the requirement to be comparable to conventional Interventional Radiology devices [35]. Reducing the size of the unbent transverse radius of curvature will also reduce our turning radius [21], [22].

**Steering cables are not housed** in the current device, aside from their final attachment to the needle tip. Tissue damage may occur as a result of these cables not remaining with the needle. This may also lead inconsistent moments initiating turns. To mitigate these issues, small flexible catheter tubing can be used route the cable along the needle body.

**Out-of-plane steering in 3D** is possible with torsional bending. The current prototypes are designed to navigate in a single plane, though any plane can be chosen initially and changed by retracting the device. At present, while the only commercially available needle is also planar, this is one of the primary disadvantages of the tape spring design versus other research needles.

**Multiple turns** with the tape spring needles are possible as we have demonstrated this. It is also likely that the current force model should apply, but this has not yet been tested and modeling is left to future work.

### E. Ultrasound Guided IV as a Steerable Needle Application

Images captured during the ultrasound IV placement trials are shown in Figure 9. While the conventional needle could reach the channel (not pictured), only a short segment of the tip could be placed within the channel and small movements caused the tip to puncture through the bottom of the channel.

**Fig. 9:**
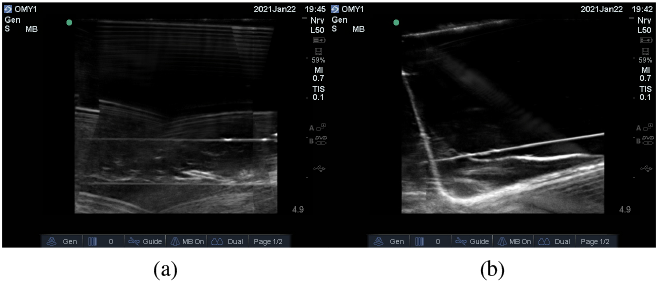
Ultrasound guided IV placement testing images. Left: 13 kPa gel with an empty 1cm diameter channel. Right: Steerable needle successfully guided inside the channel with an initial angle of approach of 90°

For this device to be a useful intraoperative tool, the needle must be compatible with one or more standard visualization modalities. The current prototype’s stainless-steel body prevents this device from being used with magnetic resonance imaging (MRI), however, computed tomography (CT) and/or ultrasound imaging are compatible. A set of ultrasound guided experiments were run in order to: demonstrate that the steerable needle design currently employed can navigate to a specific target with intuitive unaided human control; show that the needle can be accurately visualized and guided utilizing ultrasound as the only imaging modality; and show the steerable needle can be used to navigate to and through a channel (blood vessel) after having initially approached the channel at an angle too extreme for a straight needle to enter the channel. While a 90°angle is an angle that is greater than what would normally be attempted, the success of this experiment demonstrates the potential increase in maneuverability that this device could provide. Real-time visualization of the needle with ultrasound allows the user to carefully adjust the direction of the tip as the needle is moved towards the target location. In these ultrasound experiments, the tape spring steerable needle successfully navigated into a deep “blood vessel” phantom. Five successful total trials had no instances of through-and-through puncture occurring using the steerable needle. A conventional straight needle using ultrasound guidance navigated to the channel in five total trials with no success due to “through and through” puncture of the vessel at the angle of approach that was perpendicular to the model vessel and tearing of surrounding gel. While conventional IV needles may deliver a flexible catheter into the target blood vessel, this device was capable of both traveling to the target area as a conventional needle would and also traveling into the channel itself as a flexible catheter would. This highlights the ability to maintain a straight trajectory when desired and perform a small radius turn when needed. The low energy required to propagate a band down the body of the needle allowed it to easily follow the path of the “blood vessel” once inside. This application was ideal for the tape spring steerable needle as it was confined to a single plane of navigation.

## V. CONCLUSION

To maximize effectiveness, a steerable needle needs to do three things, 1) turn sharply to achieve high accuracy, 2) minimize damage to tissue and 3) maintain a useful insertion depth to reach target organs. For the first, we have shown a minimum turning radius, 3mm, lower than typical surgeon precision errors, an order of magnitude smaller than the only commercially available steerable needle, and more than twice as small as anything in the literature. We have shown the turning radius and thus angle steering is independent to tissue stiffness ranging from 7kPa to 76kPa. The small turn radius that this device is capable of performing, as well as its easy control, make it a device with large potential for clinical applications in the future.

For the other two, the tissue stresses which lead to tearing are directly related to the overall needle insertion force. The maximum depth of cut is determined by when the needle insertion fails due to column buckling. Complicating this, minimizing damage includes reducing the size of the needle, which also reduces insertion depth due to column buckling failures. This is the main failure mode of the device, which we find is best modeled as a critical bending moment under confinement. This depth is directly related to the insertion force, which is initially dominated by the cutting force. Thus, it is critical to minimize the cutting force.

Minimizing the cutting force is thus the key to optimizing all aspects of performance. Two variables that affect the cutting force, velocity and tip design, warrant further discussion. In addition, friction force is dominated by viscous friction and thus correlated with velocity. This implies decreasing insertion velocity makes failure less likely.

For this device to be of clinical utility, it must not only be able to navigate to a specific position, but it should be able to deliver specific therapies or perform certain procedures once in the correct position. Currently, payload mechanisms are being explored on how they would be integrated into the needle design. These include: integrated tissue biopsies, thermal ablation heads, and a working channel (tube from tip to base). Additionally, while insertion of the device into tissues has been explored extensively and will continue to be characterized and improved, retraction of the needle with minimal tissue damage must be explored.

